# A divide and conquer approach (DACA) to predict high fidelity structure of large multidomain protein BRWD1

**DOI:** 10.1101/2023.07.10.548240

**Authors:** Rajkrishna Mondal, Malay Mandal, Tapobrata Lahiri

## Abstract

Therapeutic importance in inhibiting Bromodomain and WD Repeat Domain containing BRWD1 against numerous human pathophysiological processes including cancers prompts prediction of a workable structure of this large protein. Here, a novel divide and conquer strategy was adopted to utilize smaller overlapping sequence-fragments of BRWD1 to further utilize their predicted structures as derived templates for prediction of complete BRWD1 structure in absence of its desired homologues in the template database. The novelty of this methodology stemmed from the requirement of templates of high sequence similarity in any comparative model based predictors whereas, the own fragments of the same target protein, BRWD1 could successfully fulfill this criteria. Additionally, the outputs of different high performing predictors including AlphaFold and RoseTTAFold were systematically integrated under the premise of Inductive Reasoning. The resultant structures were validated using existing validation parameters. Finally, a new validation paradigm was adopted to screen the best structure from the result presenting in-silico studies of known interactions of BRWD1 with various small molecules like, BD inhibitors, modified histone tails, DNA motifs and interacting proteins. The algorithm proposed in this work also paved the way for prediction of authentic structures of large size proteins.

## 1 Introduction

The bromodomain and extra-terminal (BET) family (BD2, 3, 4 and BRDT in human) are considered as attractive therapeutic targets for a wide range of malignancies because of their important role in transcription and chromatin biology[1],[2]. Dual BROMO and WD40 domain containing epigenetic reader BRWD1 is necessary for B-cell development and successful immunoglobulin recombination for antibody production[3]. It does so by genome-wide reordering of enhancer accessibility to transcription factor binding by both silencing early developmental enhancers and opening those critical for late B lymphopoiesis[4]. Interestingly, frequent occurrence of BRWD1 mutations is common in patients with idiopathic hypogammaglobulinemia. BRWD1 is associated with other human diseases e.g., Down syndrome, Immunodeficiency, and inflammatory bowel syndrome and fertility defects[5],[6],[7]. BRWD1 reads lysine acetylation in histone[4],[8] and can also bind with its unacetylated form. Most likely, BRWD1 remodels chromatin structure in association with other multimeric complexes[4],[8].

Structural insight of the BRWD1 can help to predict the unknown functional aspect of this important protein.

However, our study on abundance of disorder promoting residues in BRWD1 protein sequence reveals that the Cterminal region of the protein is highly disordered. Determination of complete BRWD1 structure by experimental method appears to be very challenging because of its large size (2320 amino acids) and rich disorderedness[9]. Artificial intelligence (AI)-powered AlphaFold 2.0 of Google DeepMind has achieved long-standing goal of accurate protein structures prediction in the 14th Critical Assessment of protein Structure Prediction (CASP14) challenge by greatly outperforming other methods[10]. Therefore, in this work, AlphaFold was first employed to generate BRWD1 protein structure that revealed a long stretch of BRWD1 C-terminal of nearly 900 residue size as coil indicating further scope of improvement to make it biologically acceptable. AlphaFold claimed that the prediction accuracy decreases for the targets where the median alignment depth is less than around 30 sequences. Additionally, the model accuracy is weaker for the proteins that have a few intra-chain or homotypic contacts[10]. Therefore, a study on the potential of all existing methods appeared to be required in spite of the individual drawbacks of each of these methods.

High-resolution (typically <1.5 Å) model can be built via comparative modelling when homologous protein templates with substantial sequence similarity are available against a target protein in PDB archive. However, the quality of the modelled structure descends with increase of target-template dissimilarity. The best performing presently available tools apart from AlphaFold, e.g., RoseTTAFold and ITASSER etc., cannot predict structure of protein of sequence size >1200 aa and >1500 respectively[11],[12],[13],[14]. I-TASSER-MTD can predict protein structure with maximum length of input sequence upto 2000 amino acids when domain definitions are not provided[15],[16]. An automated molecular graphics package, MULTIDOMAIN ASSEMBLER (MDA) of UCSF CHIMERA predicts structure of large multidomain proteins with available templates mostly non-overlapping by multiple domains assembly[17]. However, present MDA provides only the primitive structure and further refinement can be done by applying other computation methods e.g., molecular dynamics, Brownian dynamics, normal mode analysis, ab-initio modelling of linkers, constrained molecular docking, or integrative modelling incorporating additional experimental data as restraints[17]. Existing method for MDA gives only one possible conformation of interdomain linker that engaged in packing of domains and there is a very high possibility of incorporation of error in structure prediction with the increase of linker length.

To circumvent the limitations of the existing predictive models of BRWD1, especially in relation to the large size proteins, first AlphaFold was used to get the complete structure of BRWD1. Since AlphaFold predicted structure indicated further scope of improvement in the predicted structure and RoseTTAFold(CASP14 runner-up) can predict the structure of human proteins(<1200 aa) even in the absence of structural homologues11, in the follow-up steps we predicted the BRWD1 structure using a divide and conquer strategy to overcome the size-limitation using combination of other prediction methods. Briefly, structures of smaller overlapping sequence-fragments of BRWD1 were first predicted by using RoseTTAFold to produce derived templates. In the next step, the Homology application, Modeller[18],[19],[20],[21] was used to get the full length structure of BRWD1 using these smaller structures of sequence-fragments as derived templates. Significantly, this method appeared to contribute as a powerful method to correctly assemble multiple domains of BRWD1 by Modeller Package through the use of these derived templates. This resulted structure was validated and compared with AlphaFold structure using common as well as novel known interaction based validation methods under the premise of inductive reasoning in which a set of queries were prepared to know the authenticity in the architecture of the predicted domains of the structure. In-silico interaction based studies using our predicted structure of BRWD1 successfully confirmed all the presently known interacting partners, e.g., small molecule inhibitors, modified histone 3 peptides, SMARCA4, and GAGA DNA motif. Finally the structure that maximally conformed to the known interactions was chosen as the best one in our study following its corroboration with the usual validation methods. Furthermore, this structure was employed to determine yet unknown interactions of BRWD1 with components of Cohesin complex predicting it’s probable role in dynamic 3D genomic organization.

## 2 METHODS

The general algorithmic formalism for implementation of DACA (Divide and Conquer Approach) was targeted to supply derived templates for the target protein in absence of any suitable template or fold within the template or fold dataset. The specialty and novelty of this derived templates was that they were built from the own part of the target protein where first the structures of overlapping fragments of the target protein were resolved to use them further as templates in a multi-template modelling. A three level tree representation of this algorithm was illustrated in Fig. 1.

**Fig. 1.**
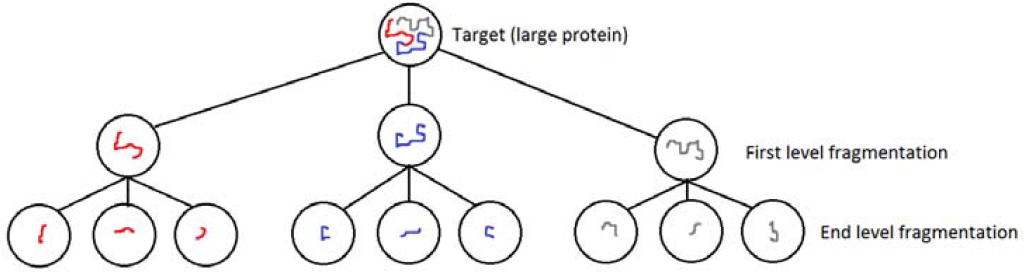
Tree representation of DACA algorithm

The proposed algorithm first needs inventory of the following knowledge-items:

i. The problem of getting an atomic level target structure through a comparative model in absence of suitable homologues was well explained by AlphaFold group [10]. This algorithm attempted to resolve it by introducing a concept of derived template which is the part or fragment of the self (i.e., the target itself).
ii. The work of Rost (1999) [22] and later on its explanation by Krieger et al (2003) [23] shows that it is not only the sequence similarity between the target and the template but also the size of the matching site that dictates the model building.
iii. Multiple templates are required for alignment correction to choose the best physicochemically viable structure out of those offered by the different templates [24].
iv. A sequence similarity of more than 75% between the target and the template theoretically guarantees building of a very accurate model structure of the target protein [23].
v. In case of using multiple templates alignments to build a model, the increase of number of templates through MSA (multiple sequence alignment) pose accumulation of errors for distant templates (i.e., the templates of less sequence similarities with the target). Heuristically in our ongoing work, we found that optimally the number of such templates should be limited to 3.

The proposed algorithm:

Thus the knowledge-items i) to v) as mentioned above, necessitates for us to consider an algorithmic formalism as described below:

Step1: Find the best probable sequence length of a protein from the database of well validated theoretically predicted protein structures (preferably those from PDB site only). We can easily get it from the sequence length distribution of these proteins. Say, this length is L.

Step2: A protein of length S (i.e., the number of residues within it) should be treated through the following recursive formalism considering K overlapping fragments at the beginning whereas, in our work the value of K is chosen as 3 heuristically:

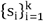

Step3: If length of si • L stop; otherwise, repeat step2 considering interim target sequence as si. Thus the abovedescribed recursion formula will build a tree structure where the furthest nodes should serve as the derived-templates of their parent nodes to build their structures, the next level parent nodes will serve as templates to give the structure of their parents, and so on to finally give the structure of the actual target protein as shown in Fig. 1.

In our present work although we found the first level fragmentation worked very well, the general algorithmic formula following the knowledge-items described above should theoretically produce a very good structure even after accumulation of errors for each recursive step since as already described, error generated at each of these recursive steps is negligible.

The second part of the algorithm to overcome limited validation by existing structure validation metrics:

The novel validation method proposed and implemented in this work although requires the information on the known interacting partners of the protein, for most of the proteins of unknown structures it is already available.

The algorithmic steps for selection of best structure by Interaction Based Validation (IBV):

Step1: List the known interacting partners of the target, e.g., ligands, other proteins, nucleic acids, etc.

Step2: For each of the models list the results of interaction as obtained through the study of in-silico interactions using docking protocols.

Step3: Select the model which best conforms with the known result of interactions.

Moreover, this work has shown the way of not only predicting a high fidelity structure but also to test its reliability at par with that determined through an experimental method.

### 2.1 Retrieval of protein sequence and its segmentation

Sequence of human BRWD1 protein (Primary accession: Q9NSI6), mouse BRWD1(Primary accession: Q921C3) and NIPBL(Primary accession: Q6KCD5) were retrieved from uniprot database and presence of putative conserved domain was analyzed by Interpro tool[25]. Full length protein sequence was fragmented into three segments, S1, S2 and S3 each one containing 1200 aa (maximum limit of RoseTTAFold tool) with the following descriptions (followed from the description of the sequence-wise domains obtained from Interpro):

S1: N-terminal segment (1 to 1200 aa) contains WD repeats domain,

S2: C-terminal segment (1121 to 2320 aa) containing BROMO domain and,

S3: middle segment containing BROMO domain (601 to 1800 aa) that overlaps with first two segments. Similarly mouse BRWD1 and NIPBL proteins were also fragmented into three segments, S1, S2 and S3 each one containing 1200 aa (Supplementary Fig. 1,2).

Overlapping segments were considered because of their potential to provide multiple templates to the Modeller for the prediction of quality structure by effectively contributing in the alignment correction step of homology modeling [24].

### 2.2 3D structure prediction of segments to be utilized as derived templates

Following the divide and conquer approach, three dimensional structure of individual protein segments were predicted using RoseTTAfold tool[11]. Total five structures resulted against every target segment were used for further work. The first and second models against each of the segments were found to score better in their structure validation as measured using SAVES v6.0. Also, in this case, the second structural model for each of the segments was found to be better than the first one. Therefore, out of 125 possible structural combinations, only 2 structural combinations were utilized in this work (Supplementary table 1.).

### 2.3 Prediction of full length BRWD1 protein structure by Multidomain Assembly (MDA) and refinement of structure

Structure of full length BRWD1 proteins and NIPBL were predicted using PyMOL plug-in PyMod 3[26] through the popular MODELLER package by Multidomain Assembly (MDA) using first and second combinations (Supplementary Table 1). Briefly, for a particular combination, first BRWD1 raw sequence was uploaded as target sequence and three different segments were considered as derived-templates backed by their structures as predicted by the method described in earlier section. It is followed by the target-template sequence alignment by inbuilt Clustal Omega tool of PyMod 3. Next, homology modelling was performed using MODELLER tool. Two structures resulted from each of the combinations were evaluated by validation tool offered by SAVES v6.0 (https://saves.mbi.ucla.edu/) and the best structure was refined further using Modrefiner tool[27].

### 2.4 Structure validation using existing methods

Full length BRWD1 structures retrieved from AlphaFold database and predicted by the present work were evaluated by different structure validation tools offered by SAVES v6.0 (https://saves.mbi.ucla.edu/) and SWISS-MODEL (https://swissmodel.expasy.org/assess).

### 2.5 Interaction based structure validation of BRWD1 using bromodomain inhibitors and histone 3

Two human histone 3 derived structure from PDB i.e., H3K9Ac (PDB ID: 5MR8, Chain:C) and H3K9AcS10PK14Ac (PDB ID: 5NND, Chain: E)[28], three bromodomain inhibitors i.e., JQ1[2], 2-(6-amino-5-(piperazin-1-yl)pyridazin-3-yl)phenol and tert-butyl 4-[3-amino-6-(2-hydroxyphenyl)pyridazin-4-yl]piperazine-1-carboxylate [29] were used for interaction study with BRWD1 through protein-protein docking using ClusPro web server [30],[31],[32],[33] and AutoDock Vina[34]. Ligand binding energy of AutoDock resulted structures were measured using PRODIGY tool[35],[36].

### 2.6 In-silico interaction studies of BRWD1 with SMARCA4, GAGA DNA and cohesin complex

SMARCA4 protein structure was collected from AlphaFold database. Cryo-electron microscopic structure of human cohesin-NIPBL-DNA complex (PDB ID: 6WG3)[37] and structure of GAGA DNA (PDB ID: 1YUI Chain: B & C)[38] were retrieved from PDB. Docking studies of BRWD1 structures with SMARCA4 and GAGA DNA were separately performed using PatchDock tool[39],[40],[41]. ClusPro online docking server was used to dock bR_BRWD1 structure with human cohesin-NIPBL-DNA complex.

## 3 RESULTS

### 3.1 Limitations of current BRWD1 structure predictions

Sequence analysis of human BRWD1 protein (2320 aa) using Interpro domain prediction tool revealed the presence of two putative BROMO domains (BDs), positioned in the region 1155-1267aa (BD1) and 1315 to 1420aa (BD2), and a putative WD40 domain (150-545aa) as shown in Fig. 2A. However, no homologue with significant similarity (>30%) was found in PDB against the full length BRWD1. Additionally, PONDR tool (http://www.pondr.com/) predicted the presence of long C-terminal disordered region from BRWD1 sequence (data not shown). Possible limitations of BRWD1 structure determination are large size of the protein, absence of homologues in structural database and presence of disordered region in the protein. Except AlphaFold, the rest of the predictors were found to lack capability to predict structure of a protein of size >2000 residues. Only, I-TASSER-MTD can predict protein structure with maximum length of input sequence upto 3000 residues provided the domain information is given which is not the case for the protein, BRWD1. Dissection of BRWD1 structure retrieved from AlphaFold (AF_BRWD1) database showed the presence of WD40 and BROMO domains that matched with prediction by the InterPro tool. In AF_BRWD1 structure, BD1 and WD40 domain were found embedded (Fig 2B, C), although BD1 and WD40 domains are preferred to be partially or fully exposed as this domains interact with other bio molecules[42]. As an evidence, major problems with the WD40 domain of AF_BRWD1 were a poor prediction of eight-bladed •- propeller and blocked central channels (Fig. 2B). On the other hand, although it seems that BD2 domain was exposed in AF_BRWD1 structure; hydrophobic tunnel of BD2 of AF_BRWD1 was partially blocked in this structure as shown in surface feeling model (Fig. 2E). This data is in striking contrast with the fact that BD2 domain have an open hydrophobic channel in experimentally determined structure of the human BRWD1 isoform A (WDR9) (PDB ID: 3Q2E, Chain: A) (Fig 1D)[8]. Furthermore, the structure predicted by AlphaFold 2.0 was found to contain a very long (about 900 aa), double size of Tau protein which is considered as one of the longest natively unfolded proteins[43], C-terminal coil region as per PDBsum analysis (http://www.ebi.ac.uk/thornton-srv/databases/cgi-bin/pdbsum/GetPage.pl?pdbcode=index.html) of AF_BRWD1 (data not shown).This data suggest a requirement of better predictable structure of BRWD1.

**Fig. 2.**
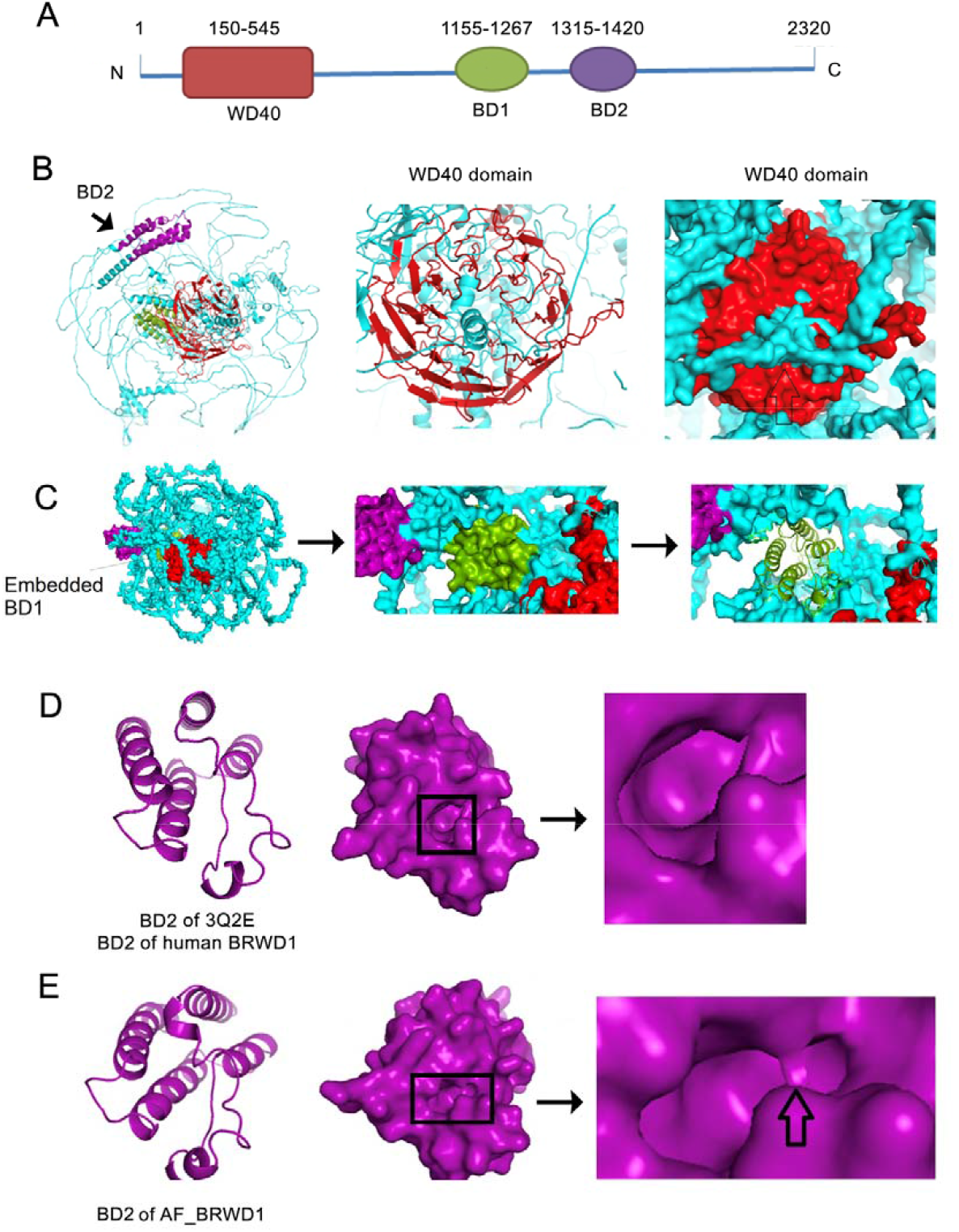
Limitations of current BRWD1 structure. A) Domain architecture of human BRWD1 protein as predicted by InterPro tool. WD40 domain (150-545aa), BROMO domain 1 (BD1 putative region 1155-1267aa) and 2(BD2 putative region 1315-1420aa) were shown in red, split pea and purple colour respectively. B) Ribbon diagram of AF_BRWD1 structure(left) showing poorly predicted •-propeller architecture of AF_BRWD1 WD40 domain(middle) and surface filling model of WD40 domain with blocked central channel (right).C)Space filling model ofAF_BRWD1 with showing embedded BD1 D-E) Ribbon diagram (left) and space filling model(middle) of the experimentally determined BD2 domain of BRWD1 (PDB ID:3Q2E) (D) and AF_BRWD1 (E). Partially blocked hydrophobic channel of BD2domain determined by of AF_BRWD1 was highlighted by vertical arrow.

### 3.2 Prediction of BRWD1 Structures by a new integrated method following divide and conquer approach

To determine a more refined structure of BRWD1, we explored a judiciously combined potential of remaining existing methods under the premise of Inductive Reasoning. To begin with, the problem of large size of BRWD1 was quite efficiently addressed by adopting two primary novel approaches. First, was a divide and conquer strategy under the paradigm of which, the large sequence of the protein in question was fragmented into some smaller overlapping segments. However, for implementation of such strategy, it was imperative to identify the information on the domains of the protein so that the fragments can be intuitively chosen. Based on the InterPro result, full length BRWD1 protein sequence was divided into 3 fragments (S1,S2 and S3) (Fig. 3A). 3D structure of each of the three segments was predicted using RoseTTAFold tool. RoseTTAFold predicted structure of segment one (S1) was found to have WD40 domain consisting eight WD40 repeats. Both S2 and S3 contains both BROMO domains (BD1and BD2), each consisting bundle of four alpha helices (Fig. 3B) that matched with prediction by the InterPro tool also. The second novel approach applied in this work was to assemble the structures obtained for all the segments using the core concept of Homology Modelling. For implementation, target sequence (complete BRWD1 protein sequence) was aligned with derived-templates using in-built Clustal Omega tool of PyMod 3. Two structures of BRWD1 were predicted using combination of derived-templates, RAW1_BRWD1 and RAW2_BRWD1 (Supplementary Table 1), by Multidomain Assembly (MDA). Resulted structures from each of the combinations were evaluated by validation tool of SAVES v6.0 (https://saves.mbi.ucla.edu/). Finally on the basis of analysis of validation scores (Supplementary Table 2), structure obtained from RAW2_BRWD1 was considered for further work and RAW1_BRWD1 was not considered. RAW2_BRWD1 structure was further optimized using Modrefiner tool and designated as ‘bR_BRWD1’ for its further utilization.

**Fig. 3.**
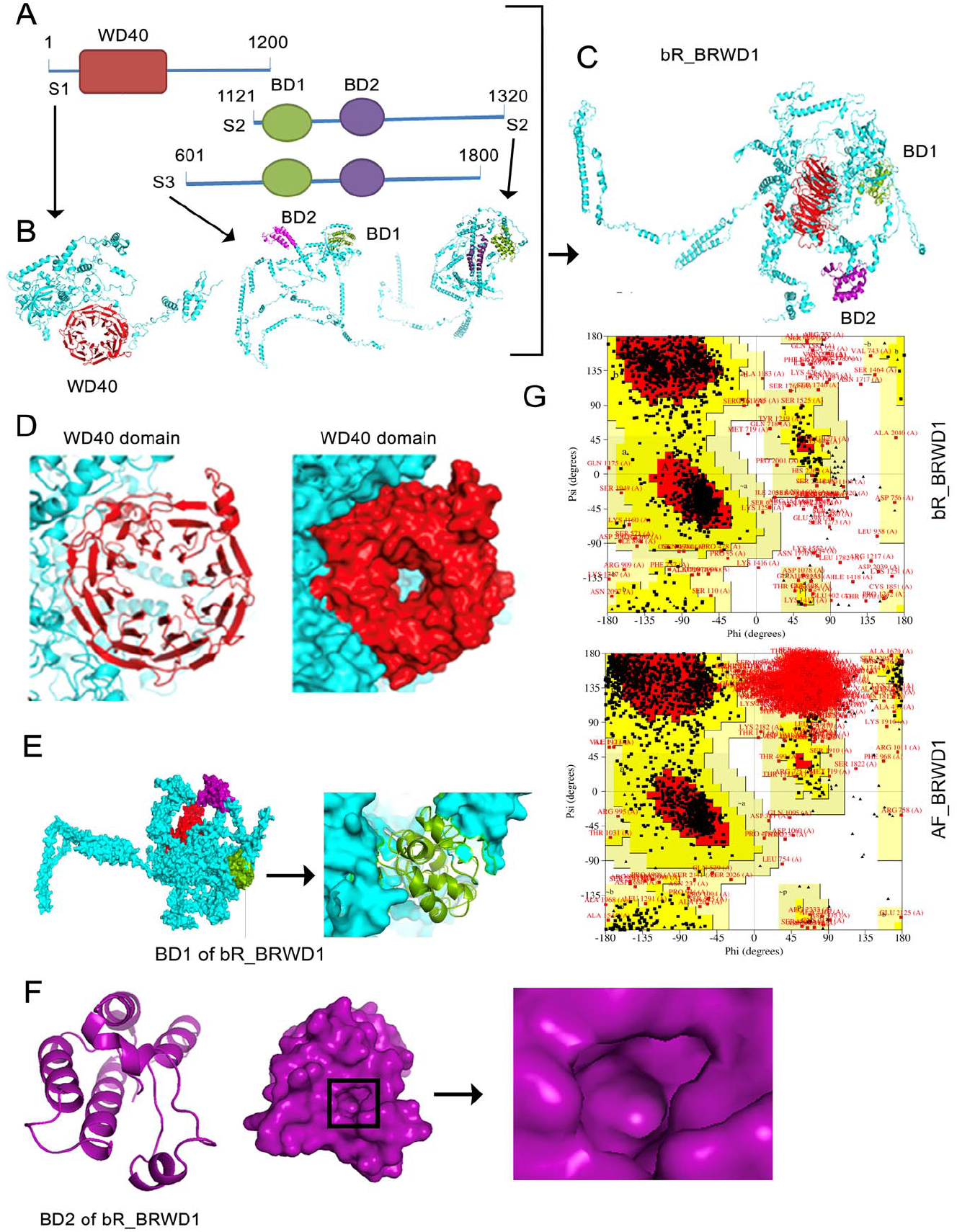
Prediction of BRWD1 Structures by a new integrated method following divide and conquer approach. A) Positional details of segments S1(1-1200aa), S2 (1121-2320aa), and S3 (601 – 1800aa) to predict fragmented structures. B) RoseTTAFold derived structure of S1. S3, and S2 segments respectively.C) Ribbon diagram of full length bR_BRWD1 structure with WD40(Red), BD1(split pea) and BD2 (purple) domains derived from S1,S2 and S3.D)Ribbon structure of 8 bladed •-propeller architecture(left) and surface filling model of WD40 domain of bR_BRWD1 (right) with opened central channel. E)Space filling model of bR_BRWD1 with exposed BD1 domain. F) Ribbon diagram (left) and space filling model (middle) of the BD2 domain of bR_BRWD1 showing opened hydrophobic channel of BD2 domain (right). G) Comparison of Ramachandran plot of bR_BRWD1 (top) and AF_BRWD1 (bottom)

Analysis of bR_BRWD1 structure revealed that both the bromodomains (BD1 and BD2) and WD40 domain were exposed as shown in Fig. 3C-E. WD40 domain of bR_BRWD1structure formed an eight bladed •-propeller with an open central channel as it should be (Fig. 3D). BD2 domain of bR_BRWD1 structure was found to have an open hydrophobic channel similar to experimentally determined structure of the BD2 of human BRWD1 isoform A (WDR9) (PDB ID: 3Q2E, Chain:A) (Fig 2F). Overall, these data suggested that bR_BRWD1 is a better predicted structure than any other available current structure of BRWD1.

### 3.3 Validation of structure prediction method

Mouse NIPBL structure (m_NIPBL) was also predicted using the above mentioned method initially by fragmenting full length protein into three segments, S1 (1-1200), S2 (601-1800) and S3 (1599-2798). However, no structure was available in AlphaFold database for the protein. m_NIPBL showed RMSD: 4.57 Å when compared with experimentally derived human NIPBL protein C-terminal domain(1163– 2804) structure (PDB ID:6WG3, Chain:E) using Pairwise Structure Alignment tool offered by PDB(https://www.rcsb.org/alignment). m_NIPBL model had a QMEANDisCo Global score of 0.58±0.05 and a QMEAN of -3.1 (Supplementary Fig. 1).

### 3.4 Validation of BRWD1 structure using common validation parameters

The BRWD1 structure (bR_BRWD1) predicted by MDA and the structure retrieved from AlphaFold (AF_BRWD1) were evaluated using standard validation tool offered by SAVES v6.0 (https://saves.mbi.ucla.edu/) (Supplementary Table 2). The Ramachandran Z-score for the bR_BRWD1 and AF_BRWD1 structure as evaluated by WHAT_CHECK were -1.832 and -6.197 respectively. Ramachandran Z-score below -4.0 indicated a serious problem with the AF_BRWD1 structure (Fig. 3G, Supplementary Table 2)[44]. PROCHEK finding of low percentage of residues in Ramachandran plot core region for AF_BRWD1 structure also supported evaluation by WHAT_CHECK. Additionally, low ERRAT value (<50) indicated poor quality of the AF_BRWD1 model[45]. Mouse BRWD1 protein structure (m_bR_BRWD1) predicted by above method outperformed the structure (m_AF_BRWD1) obtained from AlphaFold database as per the standard validation tools offered by SAVES v6.0 (https://saves.mbi.ucla.edu/) and SWISSMODEL (https://swissmodel.expasy.org/assess) as shown in Supplementary Table 2 and Supplementary Fig. 2. Calculated QMEAN for the m_bR_BRWD1 and m_AF_BRWD1 were -4.55 and -14.8 respectively.

### 3.5 Validation of BRWD1 structure using in-silico study of interactions of BRWD1 with known interactive small molecule inhibitors

AutoDock Vina based Docking study of bR_BRWD1 structures with JQ1 (a thienotriazolodiazepine and bromodomain inhibitor)[2] revealed the possible H-bond formation between ligand and Pro1337 in BD2 of bR_BRWD1 as observed for structure of the BD2 of human BRWD1 isoform A (WDR9) (PDB ID: 3Q2E, Chain:A). This interaction was however missing in case of AF_BRWD1 (Fig. 4A-D). JQ1 binding to V1342 of bR_BRWD1 was more similar to ligand interaction through conserved Val87 of human BRD4 (PDB ID: 3MXF, Chain: A) (Fig. 4 A, C). Interaction study of bromodomain inhibitor 2-(6-amino-5-(piperazin-1-yl)pyridazin-3-yl)phenol (FX5)[28] with the bR_BRWD1 protein structures revealed exactly similar binding of ligand molecule through its attachment in hydrophobic tunnel of 3Q2E and BD2 of bR_BRWD1 structure (Fig. 4E-G). However, none of the docking studies resulted complex showing binding of ligand to the hydrophobic tunnel of BD2 domain of AF_BRWD1 (Fig. 4H). Additionally, involvement of conserved Y1350 and A1389 of bR_BRWD1 in interaction with FX5 was observed similar to binding of experimentally determined structure of 5th bromodomain of human polybromo-1 (PDB ID: 6ZS3, Chain: A) [28] through Tyr696 and Ala735 respectively. Docking studies of another BD inhibitor (ligand), tert-butyl 4-[3-amino-6-(2-hydroxyphenyl)pyridazin-4-yl]piperazine-1-carboxylate (QP8) with bromodomain structures showed involvement of conserved A1389 and P1337 of BRWD1 in concurrence with that observed for 5th bromodomain of human polybromo-1 (PDB ID: 6ZS4, Chain: A) [28] via A735 and P688 respectively (Fig. 4I-L). Promisingly, majority of the interacting residues involved in QP8 inhibitors binding were common for 3Q2E and bR_BRWD1 although involvement of other residues in ligand binding were observed in case of AF_BRWD1 (Fig. 4L). Above interaction studies suggested that binding of BD inhibitors to bR_BRWD1 were more similar with experimentally proven previous findings in comparison to AF_BRWD1.

**Fig. 4.**
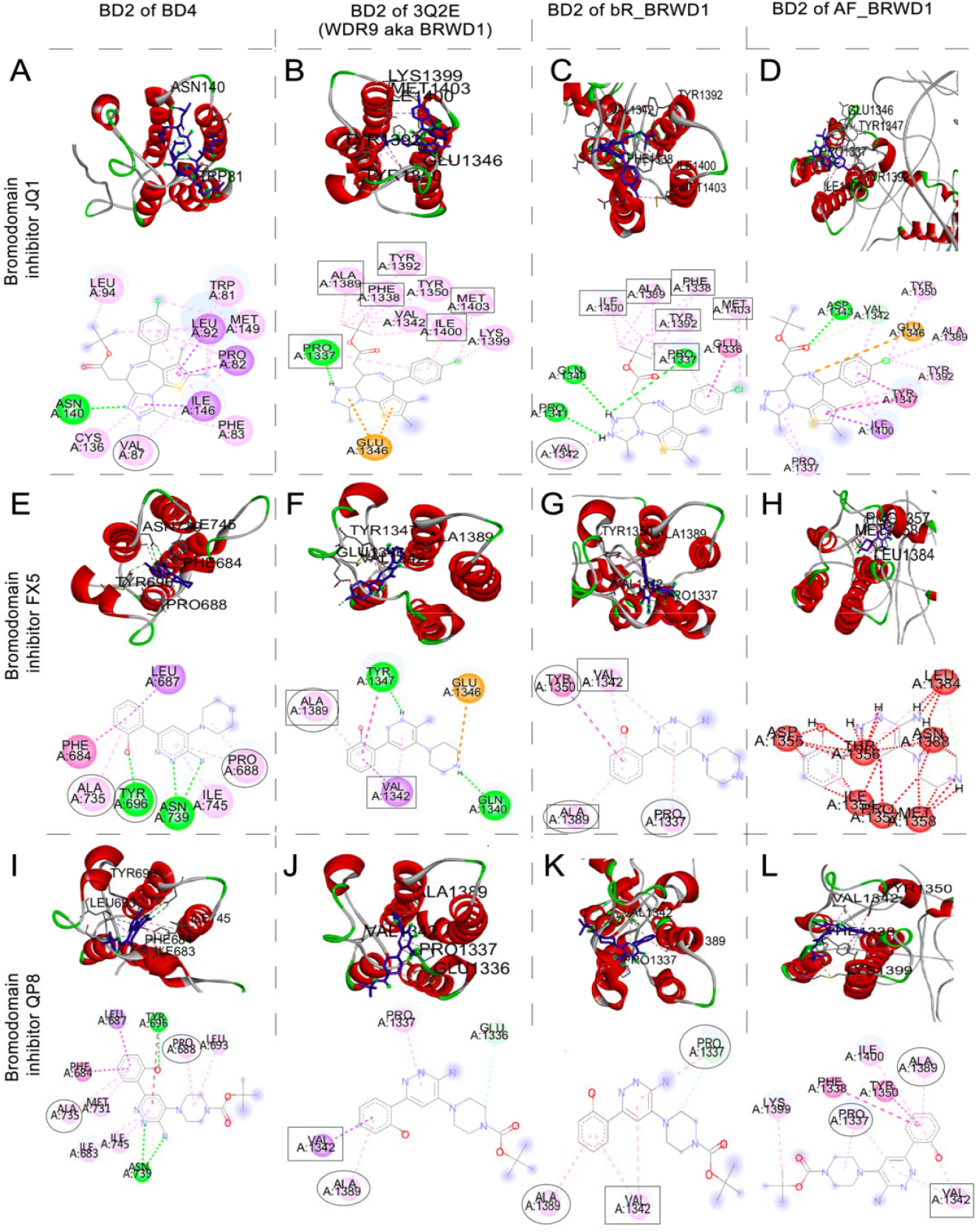
Validation of derived bR_BRWD1 structure using in-silico study of interactions of BRWD1 with known interactive small molecule bromodomain (BD) inhibitors. A-D) Key residues involved in BD inhibitorJQ1 (ligand) interaction with BD2 of human BRD4 (A), 3Q2E (B), BD2 of bR_BRWD1 (C) and BD2 of AF_BRWD1(D). E-H) Key residues involved in BDinhibitor FX5 (ligand) interaction with bromodomain of human polybromo-1 (E), 3Q2E (F), BD2 of bR_BRWD1 (G) and BD2 of AF_BRWD1(H). I-L)Key residues involved in BDinhibitor QP8 (ligand) interaction with bromodomain of human polybromo-1 (I), 3Q2E (J), BD2 of bR_BRWD1 (K) and BD2 of AF_BRWD1(L) Conserved residues of bromodomain of BRD4, 3Q2E and BRWD1 involved in interaction were encircled. Common residues of 3Q2E and BD2 of bR_BRWD1 involved in ligand interaction were labelled with rectangles. H-Bonds,Pi-sigma, amide Pi stacking and alkyl hydrophobic interactions were shown in green, purple,dark pink and light pink.

### 3.6 BRWD1 interacts with histone 3 chimeras

BRWD1 is a histone acetylation reader molecule and was shown to recognize several modifications in histone tails H3K9AcS10PK14A[3],[8]. These interactions are very important for immunoglobulin recombination via chromatin remodelling and recruitment of recombination activating proteins RAGs in lineage and developmental stage specific way[3],[46]. These interactions also take part in enhancer silencing and activation[4]. Analyses of ChIP-Seq data of BRWD1, H3K9Ac and H3S10pK14Ac (GSE63302) determined the enrichment H3K9AcS10PK14Ac in BRWD1 binding sites (Fig. 5A) and were found to co-bound genome wide. To confirm these biochemically proven interactions, bR_BRWD1 and AF_BRWD1 structures were separately docked with two H3 derived peptides i.e., H3K9Ac (PDB ID: 5MR8, Chain:C) and H3K9AcS10PK14Ac (PD**B** ID: 5NND, Chain: E)[27] using ClusPro tool to further validate the BRWD1 structures generated in this work. Th**e** logical ground behind these docking strategies were stemmed from the well-known fact that the flexible Nterminal tails of core histone proteins are protruded out from the nucleosome particles and are subjected to various post-translational modifications including acetylation [47] indicating them as the right candidate for docking. Two cluster positions of the resulted docking showed H3K9Ac interaction with first BROMO domain (BD1) of human BRWD1 (bR_BRWD1) protein (Supplementary Table 3). Docking study using ClusPro also established the interaction of H3K9AcS10PK14Ac with bR_BRWD1 through BD1 (Supplementary Table 3). This data is consistent with earlier report describing human BROMO protein BRDT interacts with nucleosomes through its first BROMO domain (BD1) but not second (BD2)1. However, AF_BRWD1 showed interaction only with H3K9AcS10PK14Ac through its second BROMO domain (Supplementary Table 3). As the modified residues of H3 derived peptides were removed during pre-docking process by ClusPro therefore the interaction of acetylated lysine and phosphorylated serine residues with BDs was not revealed.

**Fig. 5.**
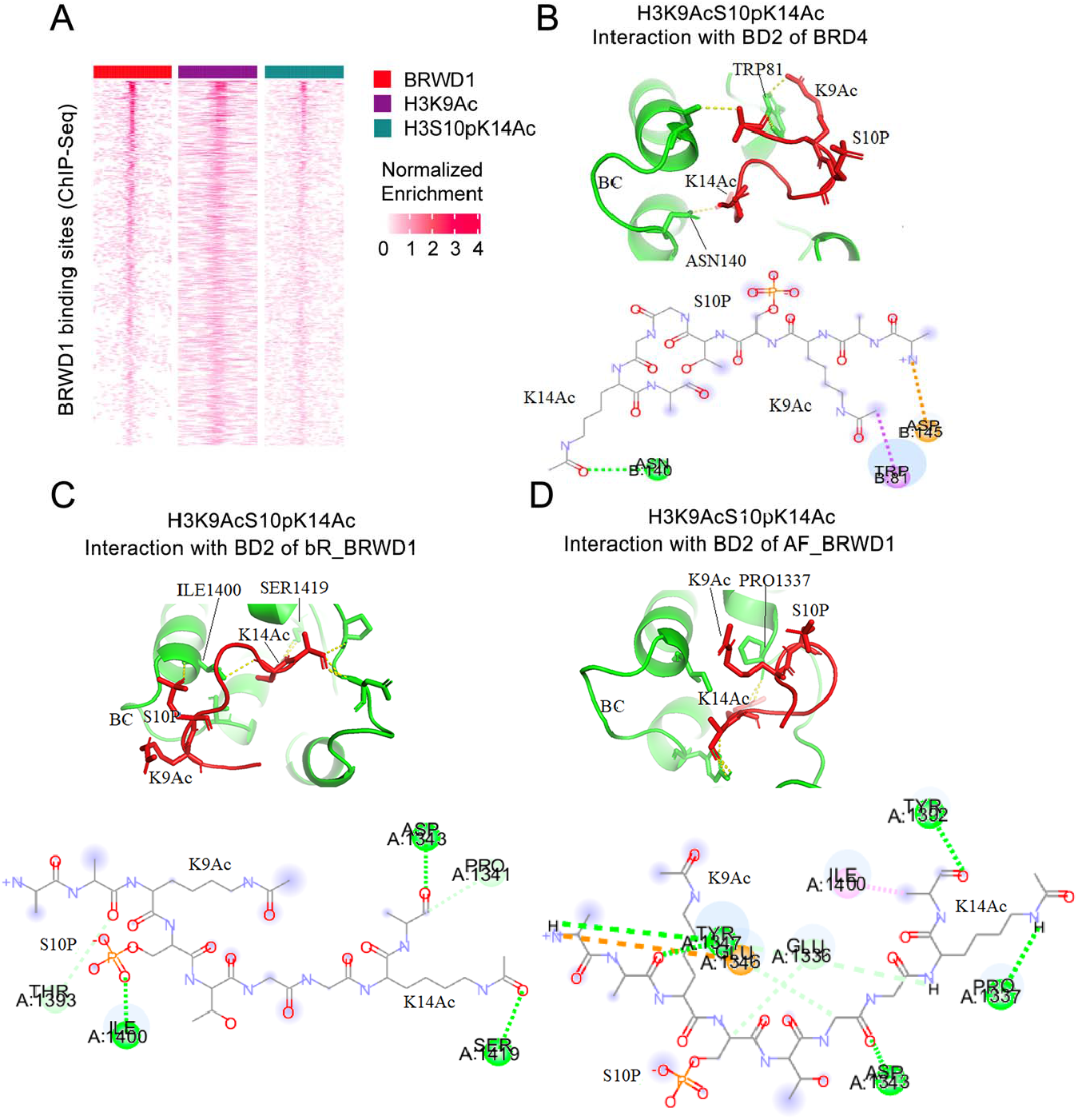
Interaction of BRWD1 with H3 histone marks H3K9AcS10PK14Ac. A)Heatmaps of tag density of BRWD1, H3K9Ac and H3S10pK14 binding along BRWD1 binding sites within ±500 base pairs of the center of the peaks binding, determined by ChIP-seqs in primarymousesmall pre-B cells progenitors B-D) Ribbon and structural diagram showing key residues of bromodomain of human BRD4 (B), BD2 of bR_BRWD1 (C) and BD2 of AF_BRWD1(D)involved in interactions with modified histone marks H3K9AcS10PK14Ac (PDB ID: 5NND, Chain: E).H-Bonds, attractive charge, Pi-sigma, amide Pi stacking and alkyl hydrophobic interactions were shown in green, brown, purple,dark pink and light pink.

BC loop asparagines of bromodomains including Asn140 of first bromodomain of human BRD4 (PDB ID: 5NND, Chain: B) were found to play critical role in interaction with acetylated lysine of histone 3 /4 proteins (Fig. 5B). . However, multiple sequence alignment result (Supplementary Fig. 3) showed that BD2 of human BRWD1 lacks conserved Asn in the preferred position. H3K9AcS10PK14Ac (PDB ID: 5NND, Chain: E) was docked with bR_BRWD1 and AF_BRWD1 structure separately targeting BD2 domain of the respective structure (Fig. 5B-D). BD2 domain of bR_BRWD1 structure showed interaction with K14Ac and S10P through Ser1419 and Ile1400 respectively as shown in Fig. 5C. However, BD2 domain of AF_BRWD1 structure read only K14Ac modification through Pro1337 (Fig. 5D). Above interaction studies revealed that binding of bR-BRWD1 to histone modifications via its BDs were more similar with experimentally proven previous findings.

### 3.7 In-silico interaction studies of BRWD1 with GAGA DNA motif

Earlier report based on ChIp-seq experiments using mouse progenitor B cells revealed that BRWD1 is recruited in GAGA DNA motifs in combination with H3K9AcS10pK14Ac as shown in Fig. 6A and played a role in sliding or evicting nucleosomes from GAGA motifs by acting like a eukaryotic GAGA factor[3],[4]. DNA binding properties of BRWD1 is yet to be explored in atomic level by standard experimental methods. Analysis of complexes resulted from the docking study of bR_BRWD1 structures with 11 bp DNA (PDB ID: 1YUI Chain: B & C)[37] containing GAGA DNA motif showed that R258 and T259 residues of bR_BRWD1 WD40 domain were involved in interaction with DNA as shown in Fig. 6B,C. However, WD40 domain of AF_BRWD1 did not show involvement in DNA binding (Fig. 6D). This data explained previously found observation of BRWD1 recruitment in GAGA DNA motif[4].

**Fig. 6.**
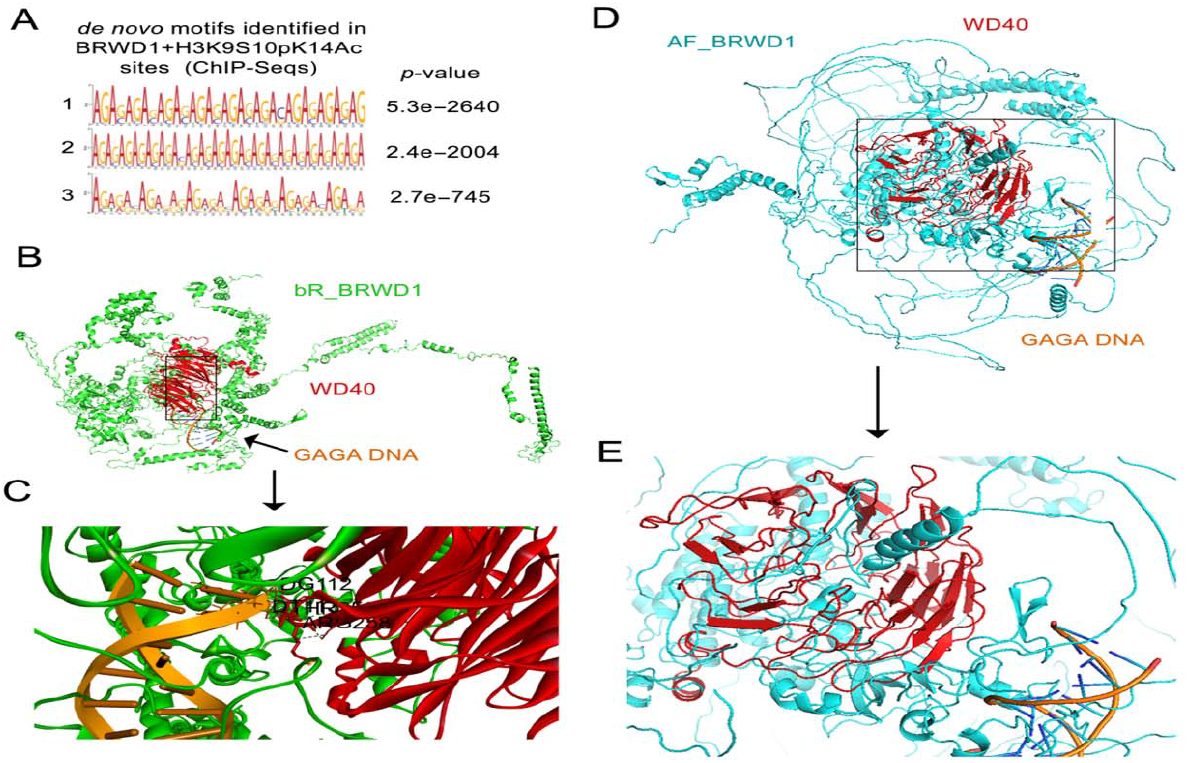
BRWD1 recognizes GAGA DNA motifs. A) de novoDNA motifs identified in the common binding sites of BRWD1, H3K9Ac and H3S10pK14Ac (determined by corresponding ChIP-seqs). B-E)Docking study of 11 bp GAGA DNA (orange) (PDB ID: 1YUI Chain: B & C) with bR_BRWD1(green) (B,C) and AF_BRWD1(cyan) (D,E) separately using PatchDock. B) & D) Ribbon diagram of bR_BRWD1-DNA complex and AF_BRWD1-DNA complex respectively C) and E) close-up look of the interaction focusing on WD40 domain (red).

### 3.8 BRWD1 interacts with SMARCA4

SMARCA4 (BRG1) is a component of SWI /SNF chromatin remodelling complexes which changes chromatin structure by altering DNA-histone contacts within a nucleosome in an ATP-dependent manner by carrying out key enzymatic activities, Co-immunoprecipitation of BRG1 with BRWD1 had established the interaction of BRWD1 with BRG1 probably via it’s WD40 repeat domain region23. Therefore, for further validation of our model structure we studied bR_BRWD1 and BRG1 interaction using PatchDocktool. Docking analyses showed that binding of AF_BRG1 (the best reliable BRG1 structure predicted by AlphaFold) with bR_BRWD1 was more stronger in comparison to its binding with AF_BRWD1 structure as found on the basis of binding energy and dissociation constant evaluated by PRODIGY for the top energetically favoured PatchDock resulted complexes (Supplementary Fig. 4). We also computed RW potential using the application calRW which was a distance-dependent atomic potential for protein structure modelling and structure decoy recognition[48]. Computation of RW potential of PatchDock resulted complexes revealed that the protein-protein complex formed by BRG1 and bR_BRWD1 (RW potential: -586461.467822 kcal /mol) was more energetically favourable compare to that formed by Brg1 and AF_BRWD1 (RW potential: - 543244.393342 kcal /mol). As expected residues of WD40 domain of bR_BRWD1 (region 150-545), viz. Y238,E239,A282,E306, and E359 interacted with BRG1 in bR_BRWD1-AF_BRG1 complex (Supplementary Fig. 4 A-D). However, direct interaction of AF_BRWD1 WD repeats with AF_BRG1 was not observed in PatchDock resulted complex (Supplementary Fig. 4 E-H). This data suggested that our model structure bR_BRWD1 is able to show the direct interaction with SMARCA4 as evidenced by immunoprecipitation experiment performed by earlier study[42].

### 3.9 Binding of predicted BRWD1 structure with components of Cohesin Complexes

Since our predicted model for BRWD1 outperformed all current models for BRWD1 structure in establishing all known interactions with small bromodomain inhibitors, like, modified histone components and GAGA DNA motif and BRG1, we explored further to predict the interactions of BRWD1 with other chromatin, and genome organizing factors. BRWD1 was shown to orchestrate enhancer landscapes by long distance genomic interactions that require 3D genomic interactions[4]. Long distance genomic interactions in mammalian cells are orchestrated by structural loops formed by loop extrusion and enhancer–promoter interactions which play a central role in gene expression and diseases[49],[50]. These dynamic genomic interactions were executed by cohesin complex which are formed by a cohesin ring composed of SMC1A, SMC3, RAD21, and STAG1 /2. Long polypeptides of SMCs fold back on themselves to form a coiled-coil domain with a hinge domain at one end and an ATPase domain at the other. The N- and C-terminus of RAD21 interact with SMC3 and SMC1A respectively. The STAG1 / 2 subunit interacts with RAD21.The NIPBL loads cohesin onto DNA to elongate loops by extrusion, whereas WAPL release cohesin from chromosomes[51],[52],[53]. Therefore, we extended our docking studies to predict if component(s) of cohesin complex interact with BRWD1. To perform this we took advantage of the recently published model of cohesin-NIPBL-DNA complex (PDB ID: 6WG3)[36] (Supplementary Fig. 5A). Strikingly, our docking studies of BRWD1 with cohesin-NIPBL-DNA complex revealed its possible interaction with SMC1A, SMC3, NIPBL and DNA through the C-terminal region (2170-2320 aa) (Supplementary Fig. 6). To cross verify interaction of BRWD1 with cohesin subunit(s) and NIPBL, we extrapolated our docking studies of bR_BRWD1 with individual subunit of cohesin-NIPBL-DNA complex (PDB ID: 6WG3). SMC1A, SMC3 and NIPBL individually showed interaction with BRWD1 (Supplementary Fig. 5B-D and Supplementary Fig. 7). Key residues of BRWD1 involved in interaction of SMC1 were D2196, R2197, N2318, A2319, and Q2320. Involvement of V2214, Q2215, P2217, S2218, E2219 and T2220 were observed for BRWD1 association with SMC3. Interaction of cohesin loader NIPBL with BRWD1 was established by the R2232, E2298, M2306, L2309, R2310, F2312, R2313, and E2317. However no noticeable interactions were obtained for RAD21 and STAG1. BRWD1 showed another type of DNA binding through its C-terminal region containing lysine and arginine residues, viz. R2179, K2180, R2195, R2204, K2222, K2224, K2229, R2304, K2305, and K2316. These observations offered a functionally testable model describing how did BRWD1 bind to DNA first and then recruit cohesin via SMC1 /SMC3 and /or form a dynamic active cohesin complex via NIPBL that was absolutely important for loop extrusion and long range genomic interactions (Supplementary Fig. 5 E).

## 4 DISCUSSION

In spite of the remarkable performances by different computational methods shown in CASP 14 competition, structure prediction of large proteins (>2000 aa) with minimum significant sequence similarity with homologues or folds still remains a challenging task. From the novel Inductive Reasoning based approach proposed in this work, it is evident that systematic integration of potentials of various structure prediction methods can give a biochemically acceptable structure better than that produced by Individually high performing structure predictive models including AlphaFold in a situation where no significant homologues or folds are available for putative domain of the protein and domains are connected by a long linker region. Our novel approach gives a BRWD1 structure that is better through its evaluation by standard structure validation parameters, visual inspection and strong authentication through novel interaction based analysis in comparison to that obtained from AlphaFold.

Overall, this work showed a novel way for deducing large protein structures under the limitations of existing predictive models including AlphaFold by augmenting the scope of existing protein structure prediction models. Initially, although the work stemmed from the intention of getting the structure of the pathophysiologically very important protein, BRWD1, it was ended up with its contribution towards resolving this problem through devising three novel strategic steps following Inductive Reasoning through listing a set of queries and corresponding probable answers for them. First query was: how to overcome the protein size limitations of existing prediction methods? Its answer brings the solution through the adoption of a “divide and conquer principle” under which paradigm the target sequence was fragmented into multiple segments. However in contrast to the common practice, the overlapping fragments were considered for doubly correct the alignment error in Modeller. In this step, the sequence-segments along with their independently predicted structures were intuitively utilized as derived-templates to get the complete structure of the large target protein for which there were no significantly acceptable templates in the usual PDB or fold database. In the next step, to overcome the limitations of common validation parameters giving contradictory indications[54], the query was: how to confirm the authenticity of the structures produced by all the different methods employed in this work? In this regard, the results of interactions of the predicted structures with the experimentally known interactive agents were also used as an interaction based validation step for them. Through the method explained and implemented in this work, a BRWD1 structure with quite high fidelity could be obtained in comparison to that obtained from the sole application of AlphaFold as validated through standard validation parameters. Furthermore, as reported above, the best structure for BRWD1 selected through the integrative approach adopted in this work was found to be highly authentic and outperformed all the individually high performing predictors including AlphaFold. This structure showed better interactions with all the interactive agents recruited in this work that corroborating well with the published reports. Similarly, mouse NIPBL structure(m_NIPBL) was predicted by above mentioned method and that showed considerable similarity with experimentally derived human NIPBL protein C-terminal domain(1163–2804) structure (PDB ID:6WG3, Chain:E). As for future implication, this novel approach may be utilized for deciphering biologically acceptable structures of large proteins competitively with AlphaFold.

Furthermore, the importance of this work to offer a high fidelity structure of BRWD1 can be understood by the recent findings on predicting its new roles in pathophysiological processes. BRWD1 apparently lacks identifiable catalytic domains. However, it is associated with nucleosome positioning probably via coordinating the activities of other chromatin regulators[3],[8]. BRWD1 can recruit the ATP-dependent chromatin-remodelling protein SMARCA4 (also known as BRG1)[42], the catalytic subunit of the mammalian SWI /SNF complex. In vivo, SMARCA4 binds immunoglobulin gene segments when they are accessible to RAG1 or RAG2 and is required for recombination[55],[56]. Docking study of bR_ BRWD1 with AF_BRG1 proved its interaction with SMARCA4 through WD40 domain as predicted by previous report. Previous studies proved DNA binding feature of BRWD1; although, structural insight of DNA-BRWD1 interaction at molecular level was yet to be explored experimentally. Our docking studies suggested binding of bR_BRWD1with GAGA part of DNA happened though its WD40 domain. Recently, our unpublished observations implied that BRWD1 orchestrates 3D genomic architecture by formation of active and dynamic cohesin complex for loop extrusion. ClusPro based docking study of bR_BRWD1 with human cohesin-NIPBL-DNA complex revealed possible interaction of BRWD1 with multiple cohesin subunit SMC31 and SMC3 in addition with cohesin loader, NIPBL and DNA through its C-terminal region. Combining two different types of DNA binding events of BRWD1 through the involvement of two different regions here we proposed a dynamic model describing how BRWD1 mediated recruitment of cohesin complex is feasible to the downstream region (Supplementary Fig. 5E). Our finding is comparable with the BRD4 interaction based stabilization of NIPBL on chromatin leading to genome folding and loop extrusion[57]. However, future experiments are necessary to validate these novel findings. Additionally, there might be other binding partners of BRWD1 which remained unidentified. Our model initiates the study to discover those unidentified partners which may have significant biological roles in cell fate decision and malignancies. Overall, considering the above observations, our predicted BRWD1 model for all the possible interactions would be a good start to understand the biology of BRWD1 function and how it works, which in future can allow us visualizing the pathway to control the diseases associated with BRWD1 and proteins related to it.

## 5 CONCLUSIONS

In summary, structure of a therapeutically important protein, BRWD1 having large size and multiple domains was obtained using a novel divide and conquer based strategy under the premise of inductive reasoning through integrating various predictors including AlphaFold. However, due to the well explained limitations of existing validation protocols to confirm acceptability of a predicted structure, in this work, a novel validation method was introduced to select the best performing structural model through in-silico studies of known interactions of BRWD1 with various small molecules, proteins and nucleic acids. BRWD1 structure selected through this stringent filtering was found to outperform current individual predictors in successfully explaining its functions. The proposed algorithm appears to resolve the problem of predicting and conclusively validating a large size protein structure.

## 6 ACKNOWLEDGEMENTS

The authors gratefully acknowledge the resources and infrastructures provided by the Department of Applied Sciences of Indian Institute of Information Technology, Allahabad to carry out this work.

