## supplemental Files for "A divide and conquer approach (DACA) to predict high fidelity structure of large multidomain protein BRWD1"

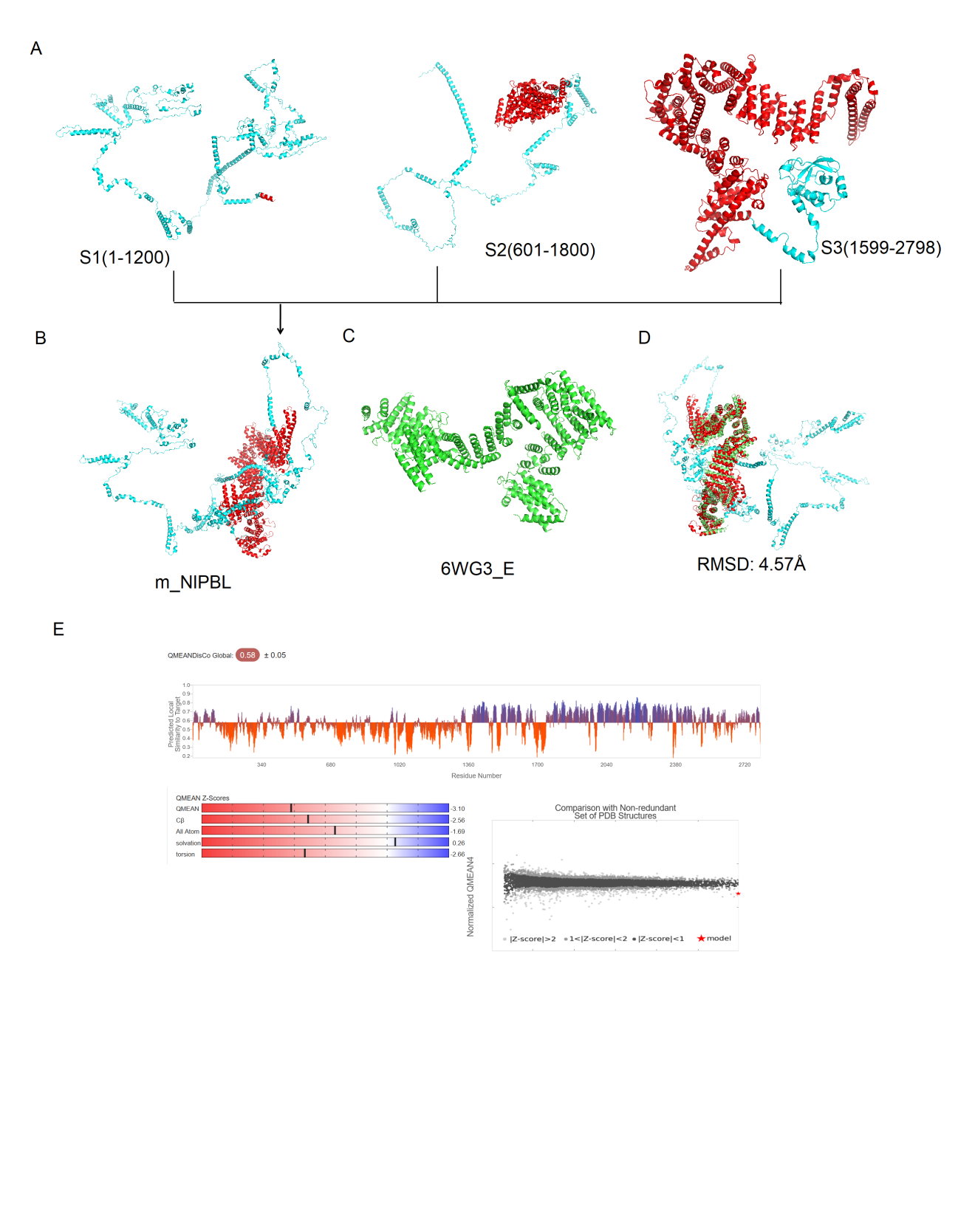

Supplementary Fig. 1. Prediction of mouse NIPBL Structure by the integrated method following divide and conquer approach. A) RoseTTAFold derived structure of S1(1-1200aa), S2 (601-1800 aa), and S3 (1599– 2798 aa) segments respectively.B) Ribbon diagram of full length m_NIPBL structure derived from S1,S2 and S3. Region of mouse NIPBL protein showed similarity with (PDB ID:6WG3 , Chain:E) heighted in red. C) Ribbon diagram of Cryo-EM structure of human NIPBL C-terminal domain (green). D) Pairwise structure alignment of m_NIPBL and 6WG3_E. E) Evaluation of the m_NIPBL structures by SWISS-MODEL assessment tool.

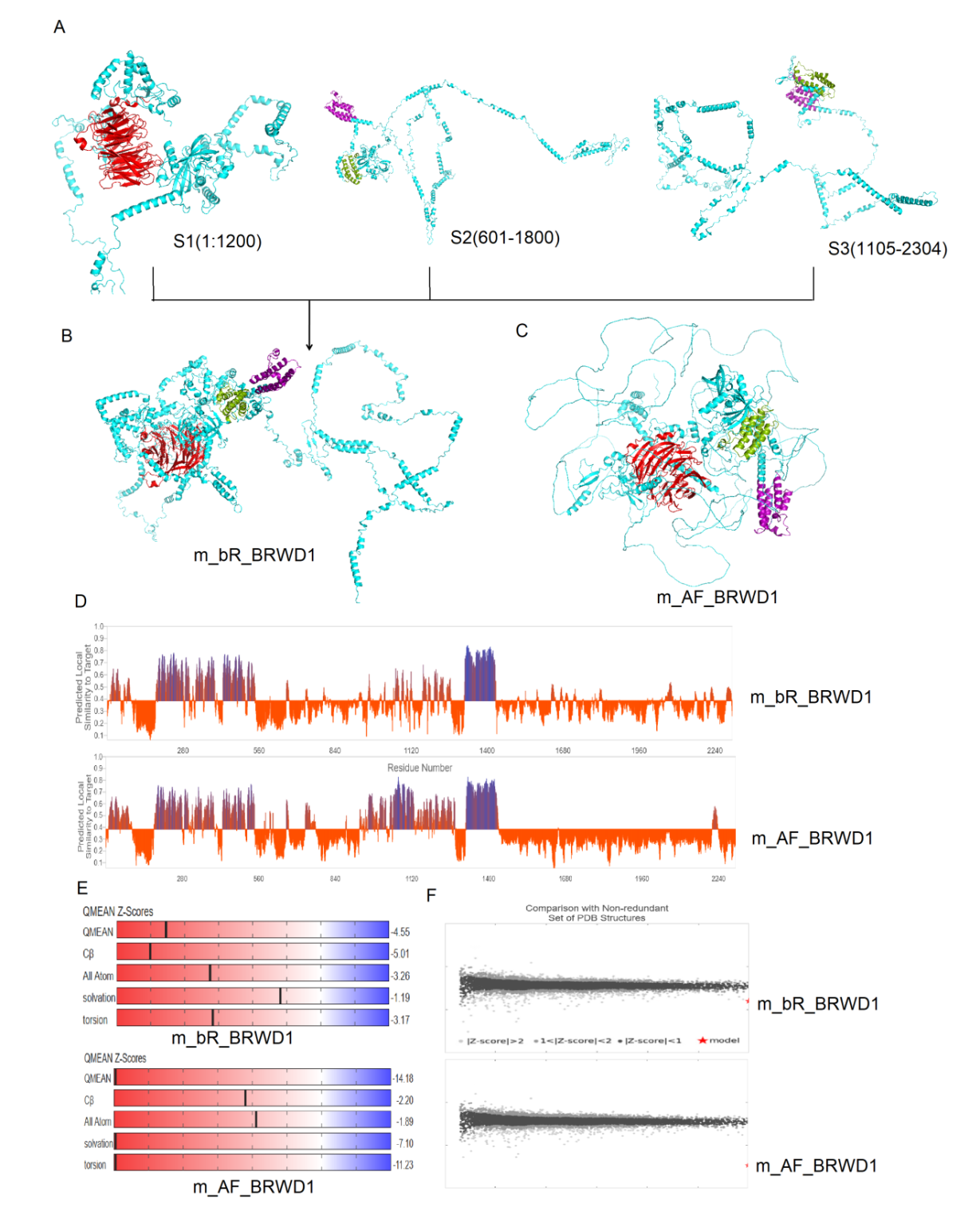

Supplementary Fig. 2. Prediction of mouse BRWD1 Structures by the integrated method following divide and conquer approach. A) RoseTTAFold derived structure of S1(1-1200aa), S2 (601-1800 aa), and S3 (1105– 2304 aa) segments respectively.B) Ribbon diagram of full length m_bR_BRWD1 structure with WD40(Red), BD1(split pea) and BD2 (purple) domains derived from S1,S2 and S3. C) mouse BRWD1 structure predicted by AlphaFold. Evaluation of the structures by SWISS-MODEL assessment tool (D, E).

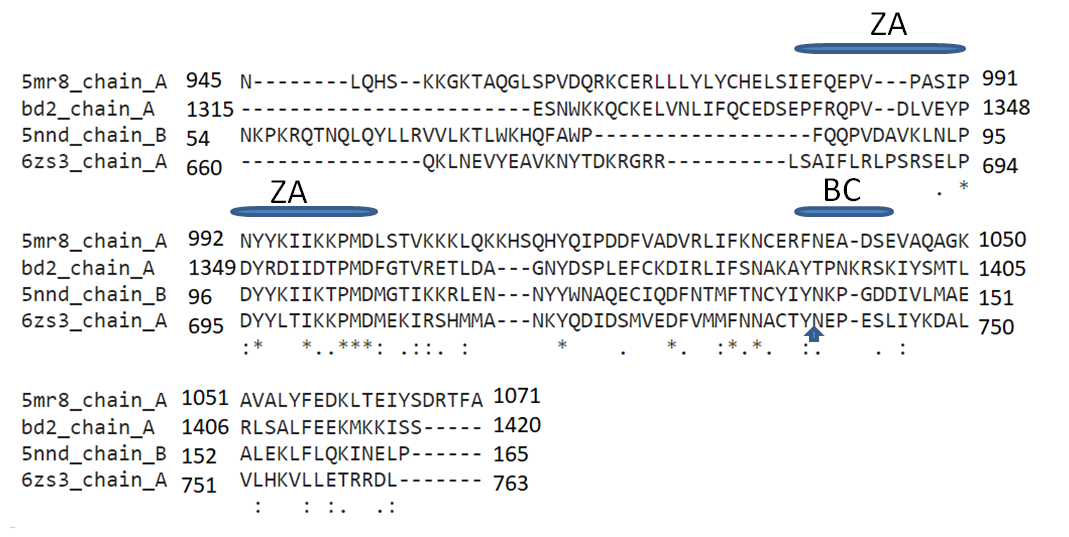

Supplementary Fig. 3. Multiple sequence alignment among bomodomain of three PDB sequences (PDB ID: 5MR8 Chain: A, 5NND Chain: B /3MXF Chain: A and 6ZS3 Chain: A/6ZS4 Chain: A) and BRD2 of human BRWD1 protein. Conserved Asn residue marked with blue vertical arrow. Positions of ZA and BC loops were highlighted by blue lines.

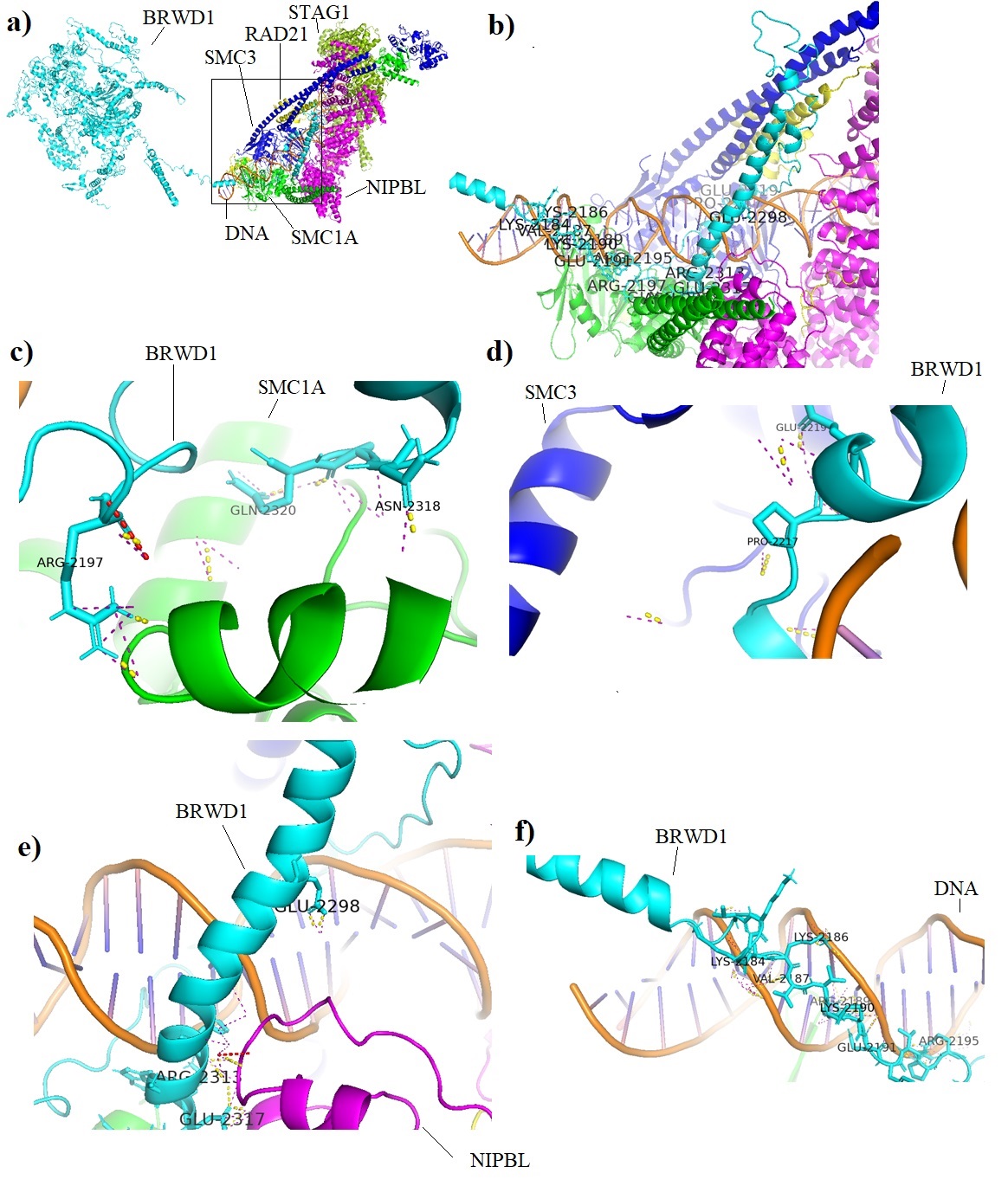

Supplementary Fig. 4. (a) Complex resulted from docking study of bR_BRWD1(cyan) with cohesin-NIPBL-DNA complex (PDB ID:6WG3) using ClusPro. b-f) Close-up look of the C-terminal end of bR_BRWD1(2170-2320) interacting with cohesion subunits, NIPBL and DNA. Key residues of bR_BRWD1 involved in interaction with SMC1A(c), SMC3(d), NIPBL(e) and DNA(f) were shown.

Supplementary Table 1. Combination of derived-templates used for prediction of full length BRWD1 protein

| Segment  Model | S1 | S2 | S3 | Combinations selected |
| --- | --- | --- | --- | --- |
| Model 1 | S11 | S21 | S31 | Combination 1 |
| Model 2 | S12 | S22 | S32 | Combination 2 |
| Model 3 | S13 | S23 | S33 | -- |
| Model 4 | S14 | S24 | S34 | -- |
| Model 5 | S15 | S25 | S35 | -- |

Supplementary Table 2. Evaluation of different structures using UCLA-DOE LAB — SAVES v6.0. structure validation parameters

| Model | ERRAT | PROVE | Ramachandran Z-score | PROCHEK | |
| --- | --- | --- | --- | --- | --- |
|  |  |  |  | Ramachandran plot | Overall G-factor |
| AF_BRWD1 | 47.7927 | Pass | -6.197 | 51.4% core 22.7% allow 8.5% gener 17.4% disall | -1.29 |
| RAW1_BRWD1 | Error | 11.2% | -0.588 | 85.7% core 10.2% allow 2.3% gener 1.8% disall | -0.49 |
| RAW2_BRWD1 | 38.8889 | 18.7 | -1.773 | 83.1% core 12.6% allow 2.6% gener 1.6% disall | -0.71 |
| bR_BRWD1 | 55.9225 | Pass | -1.832 | 82.8% core 13.0% allow 2.3% gener 1.9% disall | -0.43 |
| m_AF_BRWD1 | 83.79 | Pass | -4.830 | 60.4% core ,17.5% allow  7.4% gener ,14.7% disall | -0.85 |
| m_bR_BRWD1 | 70.123 | Pass | -0.382 | 86.4% core ,10.5% allow 1.8% gener ,1.3% disall | -0.17 |
| m_NIPBL | 81.1714 | 6.6 | 1.644 | 89.2% core 7.9% allow 1.7% gener 1.3% disall | -0.11 |

Supplementary Table 3.ClusPro results for histone 3 derived peptides docked with AF_BRWD1 and bR_BRWD1 separately.

| Protein Complex | Cluster Position | Interacting BOMO domain (Chain A) and histone peptide (Chain C/E) residues |
| --- | --- | --- |
| H3K9Ac interaction with bR_BRWD1 through BRD1 | 1 | A:ARG1162:HH22 - C:ALA1:O  A:ALA1184:H - C:THR6:O  A:ARG1213:HH11 - C:THR3:O  A:ARG1213:HH22 - C:THR3:O  A:ARG1220:HE - C:ALA1:O  A:ARG1220:HH12 - C:ALA1:O  A:GLY1187:O - C:GLN5:HE21 |
| H3K9Ac interaction with bR_BRWD1 through BRD1 | 2 | A:VAL1189:H - C:ARG8:O  A:PRO1204:O - C:ARG8:HE  A:TYR1231:OH - C:ARG8:HH21  A:PRO1204:O - C:ARG8:HH22 |
| H3K9AcS10PK14Ac interaction with bR_BRWD1 through BRD1 | 1 | A:ASN1178:HD22 - E:THR11:O  A:ASN1178:HD22 - E:GLY12:O  A:ALA1202:O - E:ALA15:H |
| H3K9AcS10PK14Ac interaction with AF_BRWD1 through BRD2 | 1 | A:ARG1339:H - E:GLY12:O  A:GLN1340:H - E:GLY12:O  A:ARG1351:HE - E:ALA15:O  A:ARG1351:HH22 - E:ALA15:O  A:PRO1337:O - E:ARG8:H  A:TYR1392:OH - E:ARG8:HH22  A:VAL1342:O - E:ALA15:H |
